# SelNeTime: a python package inferring effective population size and selection intensity from genomic time series data

**DOI:** 10.1101/2024.11.06.622284

**Authors:** Mathieu Uhl, Paul Bunel, Miguel de Navascués, Simon Boitard, Bertrand Servin

**Affiliations:** CBGP, Université de Montpellier, CIRAD, INRAE, IRD, Institut Agro Montpellier, Montpellier, France; CEFE, CNRS, Montpellier, France; GenPhySE, Université de Toulouse, INRAE, INPT, Castanet-Tolosan, France

## Abstract

Genomic samples collected from a single population over several generations provide direct access to the genetic diversity changes occurring within a specific time period. This provides information about both demographic and adaptive processes acting on the population during that period. A common approach to analyze such data is to model observed allele counts in finite samples using a Hidden Markov Model (HMM) where hidden states are true allele frequencies over time (i.e. a trajectory). The HMM framework allows one to compute the full likelihood of the data, while accounting both for the stochastic evolution of population allele frequencies along time and for the noise arising from sampling a limited number of individuals at possibly spread out generations. Several such HMM methods have been proposed so far, differing mainly in the way they model the transition probabilities of the Markov chain. Following Paris et al. (2019a), we consider here the Beta with Spikes approximation, which avoids the computational issues associated to the Wright-Fisher model while still including fixation probabilities, in contrast to other standard approximations of this model like the Gaussian or Beta distributions. To facilitate the analysis and exploitation of genomic time series data, we present an improved version of Paris et al. (2019a) ‘s approach, denoted SelNeTime, whose computation time is drastically reduced and which accurately estimates effective population size (assuming no selection) or the selection intensity at each locus (given a previously estimated value of N). We also evaluate the performance of this method in realistic situations where selection is present and both demography and selection need to be inferred. SelNeTime is implemented in a user friendly python package, which can also easily simulate genomic time series data under a user-defined evolutionary model and sampling design.

## Introduction

One limitation of most population genetic analyses is the need to use data from a single time point, the present. The genetic diversity observed in this single sample is very informative about past population history, but it integrates the effect of many evolutionary processes acting on a large time-scale, which may be hard to clearly separate. For instance, selection constraints acting on a given locus often fluctuate along time as a result of environmental changes, and such complex and changing processes are difficult to reconstruct from the observation of the resulting genetic diversity at a single recent time point. In contrast, the study of samples collected at several different generations allows one to directly measure changes in the genetic diversity of populations, informing about the evolutionary processes acting on that specific time period. Such genomic time series arise in different contexts and time scales, such as the monitoring of wild or domestic populations (Boitard et al., 2023), experimental evolution (Long et al., 2015) or ancient DNA (Fages Antoine et al., 2019) studies.

A common approach to analyze genomic time series data is to model observed allele frequencies using a Hidden Markov Model (HMM); this approach allows one to compute the full likelihood of the data, while accounting both for the stochastic evolution of population allele frequencies along time, under the combined effects of demography and selection, and for the noise arising from sampling a limited number of individuals at different generations (Bollback et al., 2008). Several such HMM methods have been proposed so far, differing mainly in the way they model the transition probabilities of the Markov chain. Indeed, while the Wright-Fisher (WF) model appears as a natural choice to model these transitions, it leads to transition matrices whose dimension is equal to the population size so it is limited in practice to small populations. Among the various approximations of the allele frequency transition distribution that have been proposed to overcome this issue (e.g. Gaussian, Beta), the Beta with Spikes (BwS) distribution (Tataru et al., 2015) was pointed out by Paris et al. (2019a) as particularly promising, because it combines a continuous (Beta) density with discrete fixation probabilities at frequencies 0 and 1. These features provide accurate estimations for a large range of demographic and selective parameters (Paris et al., 2019a), while previous methods were either impossible to run in large populations or inaccurate for some parts of the parameter space (e.g. strong selection).

However, the current implementation of this method, compareHMM (Paris et al., 2019b), considers a fixed effective population size provided by the user, assuming that this parameter is known or has been previously estimated from another software. Besides, it was initially implemented as a test tool comparing different approximations of the Wright-Fisher model in the context of genomic time series analyses, not to be used by a large community. In particular, it was not optimized to analyze very large data sets such as those typically found in the genomics era.

With the objective of improving on these several limitations, the SelNeTime python package (Uhl et al., 2025) provides a user friendly and computationally efficient implementation of Paris et al. (2019a)’s method, which estimates effective population size or selection from genomic time series. SelNeTime can also simulate such data in order to easily test the expected performance of the approach under custom sampling designs. This package is available on the python package index https://pypi.org/project/selnetime/ and on the project git repository https://forge.inrae.fr/genetic-time-series/selnetime.

## Materials and Methods

## Modelling genomic time series through an HMM

Let us consider a population for which genomic data has been collected at *n* sampling times *t*_1_, …, *t*_*n*_. For a given bi-allelic locus, we model the evolution of the (arbitrary) reference allele frequency along time using a Hidden Markov Model (HMM). We denote *X*_*i*_ the (hidden) population allele frequency at time *t*_*i*_ and *Y*_*i*_ the (observed) sample allele frequency at time *t*_*i*_. As all HMMs, this stochastic process is fully determined by three components : the initial distribution of *X*_1_, the transition distributions from *X*_*i* − 1_ to *X*_*i*_ (*i* ∈ 2, …, *n*) and the emission distributions of *Y*_*i*_ given *X*_*i*_ (*i* ∈ 1, …, *n*). Here we assume that *X*_1_ is distributed uniformly in [0, 1] and that observed frequencies result from a binomial sampling within the population, i.e.

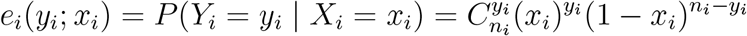

where *y*_*i*_ is the number of reference allele copies observed at time *t*_*i*_ and *n*_*i*_ the haploid sample size (i.e. total number of allele copies) at this time. Transition distributions, which make the link between observed data and the parameters of interest, are described in the next two sections.

### Modelling transitions through the Wright-Fisher model

Let us now assume that the studied population evolves under a Wright-Fisher (WF) model, i.e. with panmixia, non overlapping generations and a constant (haploid) size *N*. Under this model, allele frequencies *X*_*i*_ are discrete and take values in 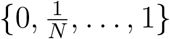. We ignore mutations due to the short time scale of interest, but model selection using a diploid codominant fitness function with selection coeficient *s*, i.e. the fitnesses of homozygote reference, heterozygote and homozygote alternative genotypes are respectively equal to 1, 1 + *s/*2 and 1 + *s*. For a single generation, this results in the transition probabilities

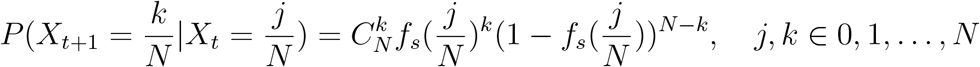

where

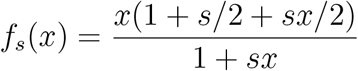

Denoting *Q*_*s,N*_ = ((*Q*_*s,N*_)_*j,k*_)_0*≤j,k≤N*_ the matrix of these single generation transition probabilities, *Q*_*s,N,i*_ the transition probabilities from *t*_*i−*1_ to *t*_*i*_ can be computed as

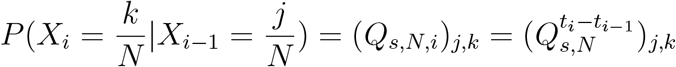

### The Beta with Spikes approximation

Because *Q*_*s,N*_ has dimension (*N* + 1) × (*N* + 1), likelihood computations under the WF model described above are only possible for small values of *N*. Following Paris et al. (2019a), SelNeTime computes (by default) this transition matrix based on a Beta with Spikes (BwS) approximation of the WF model (Tataru et al., 2015). This model assumes that the probability density of *X*_*i*_ is given by

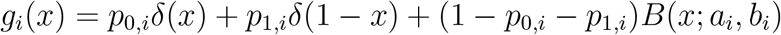

where *p*_0,*i*_ and *p*_1,*i*_ are the fixation probabilities in 0 and 1 at time *t*_*i*_, *δ*(*x*) the Dirac delta function and *B*(*x*; *a*_*i*_, *b*_*i*_) is the probability density of a beta distribution with parameters *a*_*i*_ and *b*_*i*_. While the continuous part modelled by *B*() allows one to control the size of *Q*_*s,N,i*_ using a discretization grid that is independent of *N*, accounting for the discrete fixation probabilities *p*_0,*i*_ and *p*_1,*i*_ was shown to provide better results compared to other standard continuous approximations of the WF (Paris et al., 2019a).

Following Tataru et al. (2015), the transition matrix *Q*_*s,N,i*_ from time *t*_*i−*1_ to time *t*_*i*_ is computed in three steps. 1) We compute *m*_*i*_(*N, s, x*) = *E*[*X*_*i*_ | *X*_*i−*1_ = *x*] and 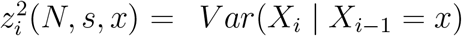 under a Wright-Fisher process, using a recursion from *t* = 1 to *t* = *t*_*i*_ *−t*_*i−*1_. For *s* = 0, this recursion relies on a large *N* Taylor expansion called Delta method (Lacerda & Seoighe, 2014). 2) Using another recursion from *t* = 1 to *t* = *t*_*i*_ − *t*_*i* −1_, we set the values *p*_0,*i*_(*N, s, x*), *p*_1,*i*_(*N, s, x*), *a*_*i*_(*N, s, x*) and *b*_*i*_(*N, s, x*) ensuring that the mean and variance of the resulting BwS distribution match the values *m*_*i*_(*N, s, x*) and 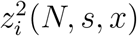 obtained at step 1. 3) The line of *Q*_*s,N,i*_ corresponding to a starting frequency of *x* is derived from the BwS distribution characterized at step 2. Detailed equations used at steps (1-2) can be found in Paris et al. (2019a).

For step 3, we introduced a technical difference with the approach of Paris et al. (2019a). In the original study, the quantities stored in *Q*_*s,N,i*_ were a mixture of probability and density values: the first and last columns represented the discrete probabilities of arriving respectively in 0 and 1, but other columns represented probability densities at arriving frequencies, which were later (numerically) integrated during likelihood computation. While more elegant mathematically, this approach occasionaly lead to numerical issues and implied that likelihood computations could not be fully vectorized, because different parts of the matrix were processed differently. In contrast, SelNeTime implementation considers that all coefficients in the matrix correspond to discrete probabilities. More precisely, for a discretization grid of (0, 1) into *n*_*x*_ bins *I*_*j*_ = [*m*_*j*_, *M*_*j*_], *j* ∈ 1, …, *n*_*x*_, *Q*_*s,N,i*_, has dimension (*n*_*x*_ + 2) × (*n*_*x*_ + 2) where *I*_0_ corresponds to an allele frequency of exactly 0 and 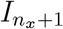 to an allele frequency of exactly 1.

Its coefficients are defined by

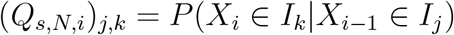

For *j* ∈ 1, …, *n*_*x*_ (i.e. when starting from a discretization bin), this probability is approximated by 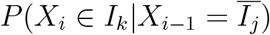, where 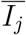 is the middle of *I*_*j*_. For *k*∈ 1, …, *n*_*x*_ (i.e. when arriving into a discretization bin), the probability is derived as *G*(*M*_*k*_) − *G*(*m*_*k*_), where *G*() is the cumulative density function of the Beta distribution obtained at step (2).

### Likelihood computation and optimization

For an initial distribution *ν*, transition matrices *Q* = (*Q*_2_, …, *Q*_*n*_) and emission distributions *e* = (*e*_1_, …, *e*_*n*_), the likelihood of observed data in a HMM can be computed using the so-called forward algorithm. This algorithm focuses on the joint probability of observations *Y*_1_ to *Y*_*i*_ and hidden state *X*_*i*_

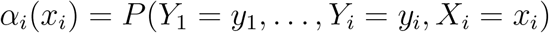

Note that the *α*_*i*_(*x*_*i*_)’s are discrete probabilities (rather than continuous measures) here, consistent with the above assumption of discretized allele frequencies. The algorithm is initiated by

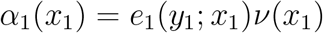

for all possible values of *x*_1_ and then updated recursively, for *i* = 2, …, *n*, by

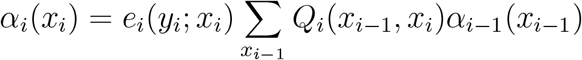

for all possible values of *x*_*i*_. The notation *Q*_*i*_(*x*_*i−*1_, *x*_*i*_) is used here for convenience to refer to the entry of *Q*_*i*_ whose row corresponds to *x*_*i−*1_ and whose column corresponds to *x*_*i*_. The likelihood is finally obtained by summing over *x*_*n*_:

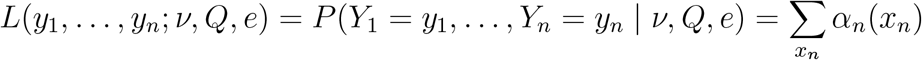

In our application, the parameters of interest are *N* and *s*, which influence Q but not *ν* and *e*. Thus, the forward algorithm provides

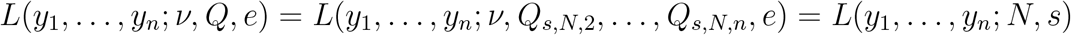

In other words, we are able to evaluate numerically the likelihood of the data for any given values of *N* and *s*. Note that this likelihood function also implicitely depends on the time intervals *t*_*i*_ *− t*_*i*− 1_, because these affect the *Q*_*s,N,i*_ matrices (see section ‘Modelling transitions through the Wright-Fisher model’). We omit this dependence here because we consider these parameters as known fixed values, but this could in principle be relaxed in applications where sample age or generation time is unknown.

In the original study of Paris et al. (2019a), *N* was assumed to be known and the likelihood was optimized over *s* (see below) to get the Maximum Likelihood Estimator (MLE) of selection intensity at the studied locus. One novelty in SelNeTime is to also estimate *N* by cumulating data from *L* loci. Indeed, *N* reflects the demography of the population so a single value applies to the whole genome. Denoting **y**_*l*_ = (*y*_*l*,1_, …, *y*_*l,n*_) the sequence of observed reference allele counts at locus *l*, the likelihood of *N* is evaluated as

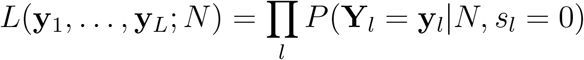

This equation is actually a composite likelihood, as it assumes that the loci are independent (which almost holds in practice if these loci are taken sufficiently far away from each other, but see the discussion). Composite likelihoods are widely used in population genetics and are theoretically unbiased (Wiuf, 2006), although they may result in too narrow (resp. too large) likelihood surfaces when loci are positively (resp. negatively) correlated. Another important assumption in the above equation is that the effect of selection at each locus can be neglected in first approximation while estimating *N*, i.e. the likelihood of *N* at each locus *l* is computed assuming *s*_*l*_ = 0; the impact of this approximation on estimation accuracy will be explored through simulations in the manuscript.

To obtain the MLE in practice, we first evaluate the likelihood on a broad grid of candidate *N* values ranging from 10 to 1,000,000, and next perform a refined search on a second grid of candidate *N* values centered around the value maximizing the likelihood in the first grid. The likelihood profile obtained from this refined search provides the MLE as well as a 95% confidence interval for *N*. Importantly, note that only integer values of *N* are explored during this two step search (besides the biological realism, several steps of the computations require integer *N* values, both for the WF and the BwS model).

Once *N* has been estimated, SelNeTime estimates the selection intensity at each locus *l* based on this value using the same approach as Paris et al. (2019a), i.e. optimizing 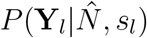 over different values of *s*_*l*_ in order to find the MLE (and confidence interval) of this parameter at each locus. As indicated in the notation, *s*_*l*_ is a locus specific parameter, because selective contraints typically differ between loci.

## Important features

### Estimation of effective population size under neutrality

Using computer simulations, we checked the accuracy of SelNeTime estimation of *N*, a task that has not been investigated by Paris et al. (2019a). We first considered simulation scenarios without selection, in order to evaluate the method in the favorable situation where its assumptions are met (except that simulations were performed under the true WF model, not the BwS approximation). Estimations of *N* in the presence of selection are investigated in a later section. As a comparison, we also estimated *N* for the same data with the method of Hui & Burt (2015), implemented in the NB package (Hui, 2014), which is based on a continuous Beta approximation of allele frequency transitions. Interestingly, both methods performed equally well for short sampling time intervals (Δ*t* = 1) but the accuracy of NB was reduced for larger time intervals (Δ*t* = 10) (Figure 1); this is consistent with the higher probability of fixation events, which are not accounted for in the method of Hui & Burt (2015), when Δ*t* increases, illustrating the advantage of the BwS approximation.

**Figure 1.**
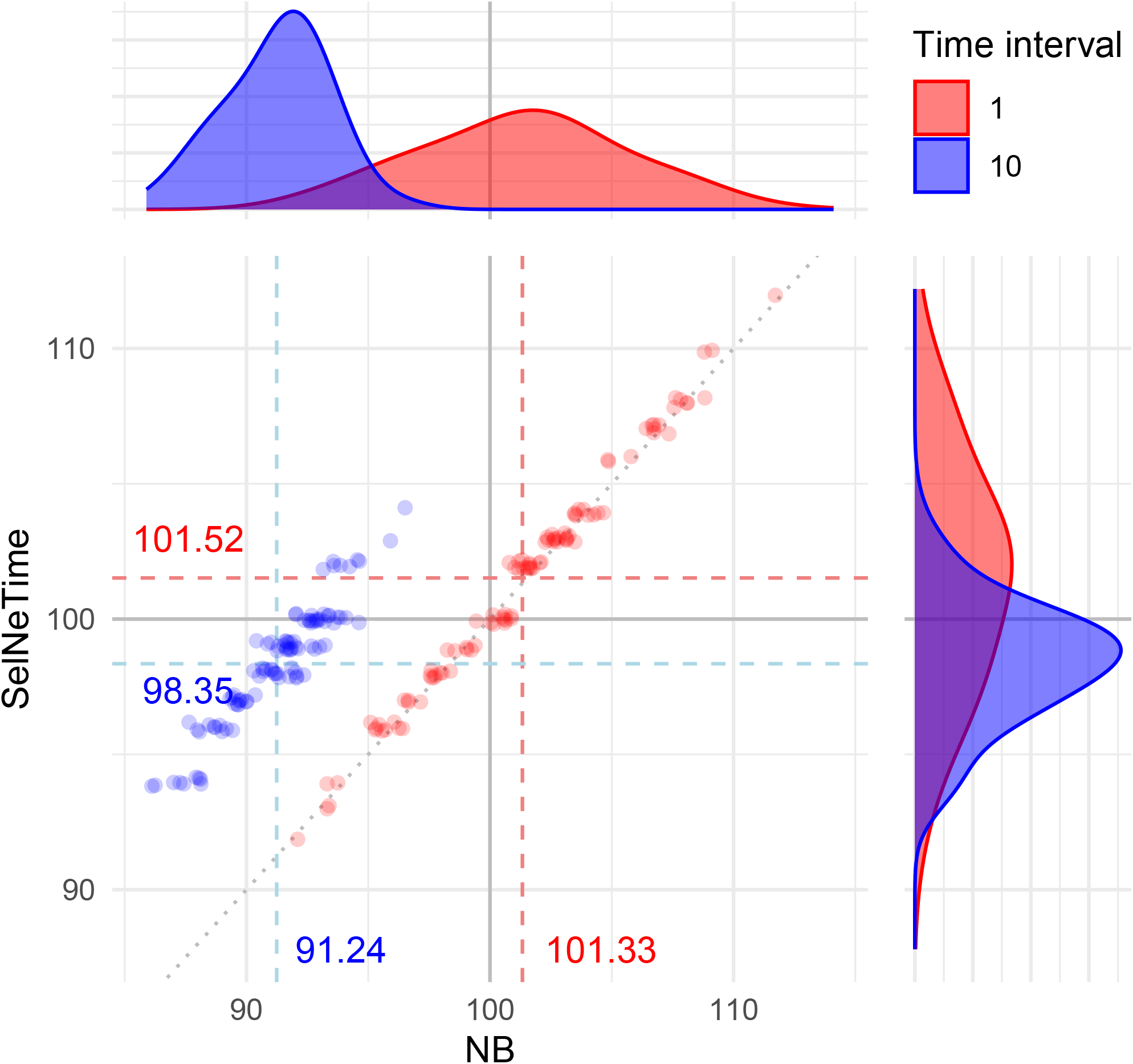
Estimation of the effective population size for 100 datasets, using SelNeTime or NB (Hui, 2014). Each dataset includes 1000 loci and 10 sampling times *t* = 1 … 10 (red) or *t* = 10, …, 100 (blue). Allele frequency trajectories at each locus were simulated independently under the Wright-Fisher model with *N* = 100, *s* = 0 and an initial allele frequency *x*_0_ = 0.5. At each time point *t*_*i*_, sample size is *n*_*i*_ = 30.

We also explored neutral scenarios with different values of *N* and of the initial allele frequencies, in order to evaluate the influence of these parameters on SelNeTime estimations. SelNeTime performed almost equally well for all choices of the initial allele frequencies, with a slight advantage for trajectories starting from *x*_0_ = 0.5 (Figure S1). This can be explained by the higher probability of fixation events when the initial allele frequency is close to 0 or 1. Indeed, while these events are accounted for in our method, the effect of approximating the true (simulated) WF distribution by the BwS distribution is stronger when the probability of fixation events increases. Estimations of *N* were also less accurate (in particular they showed a higher variance) for large populations, as expected given the more subtle effect of genetic drift in such scenarios. However, results obtained with *N* = 10, 000 (the largest value tested here) remain very good. They could also be improved by increasing the sample size at each time point, which would reduce the sampling variance and thus make the drift variance easier to capture.

### Estimation of selection intensity at a locus using the true value of *N*

We next evaluated the accuracy of SelNeTime to estimate *s*_*l*_ at each locus, assuming that *N* is known (i.e. that it has been correctly estimated). As this method is essentially a re-implementation of the BwS approach described by Paris et al. (2019a), we first compared it with the inital implementation published by these authors on a limited number of scenarios (Paris et al., 2019b). As expected, we found very similar results between the two implementations (Figure S2), which both lead to almost unbiased estimations of the selection intensity.

To provide a first hint of SelNeTime performance in more diverse scenarios, we then ran it on datasets simulated with various population sizes, selection coefficients and sampling times, with the true value of *N* still provided (Figure 2). Results showed a larger estimation variance for lower values of *N* and Δ*t*, but the mean of estimated values was generally very close to the true value of *s*. A more detailed investigation of the estimation accuracy and of the power to detect selection can be found in the study of Paris et al. (2019a), which explored these questions for a large range of parameter values and sampling designs.

**Figure 2.**
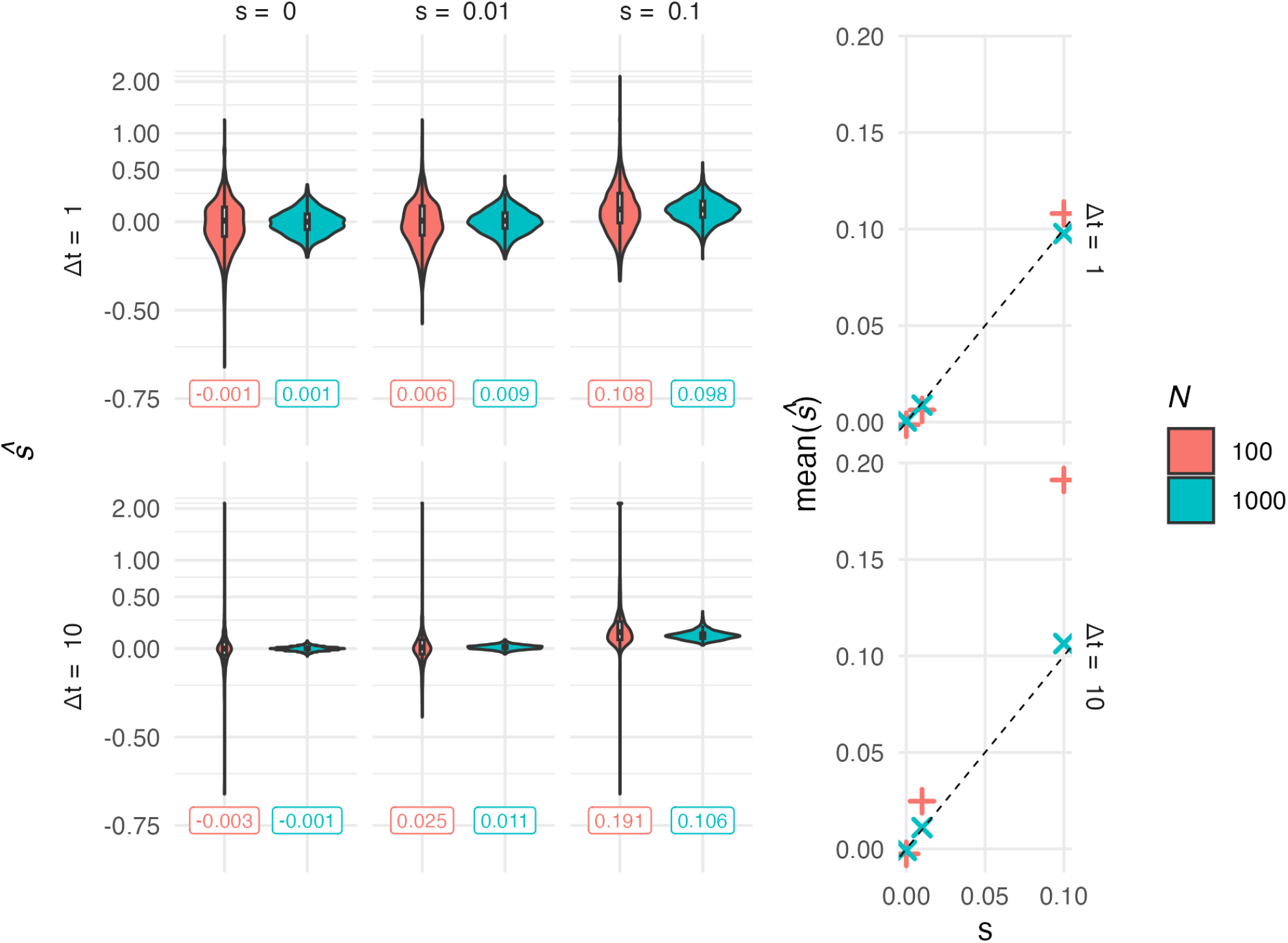
Estimation of the selection intensity using SelNeTime with the true value of *N*. For given values of *s* (columns) and *N* (colors), 1000 loci were considered. Allele frequency trajectories at each locus were simulated independently under the Wright-Fisher model, with *n* = 30 at each of 10 sampling times *t* = 1, …, 10 or *t* = 10, …, 100 (lines), and an initial allele frequency *x*_0_ = 0.5. Distribution of estimations for the 100 loci. The mean value for each simulation scenario is shown below the associated violin plot, and on the right part of the plot. A log_2_(1 + *x*) transformation was applied to the y axis of the left panel.

### Estimation of effective population size and selection intensity when both are unknown

Finally, we considered the more realistic situation where some loci are under selection and both *N* and the *s*_*l*_ need to be estimated. In such situations, we remind that SelNeTime would first estimate *N*, assuming that all loci are neutral, and then estimate *s*_*l*_ at each locus based on the obtained value of *N*. To evaluate the accuracy of this approach, we first simulated favorable scenarios where only a minor fraction of the genome (here 1%) is under selection, following a classical assumption in population genetics. As shown in Table 1, estimations of *N* remain very accurate under these conditions, unless when both *N* and *s*_*l*_ are high where a small negative bias is observed. Mean estimations of *s*_*l*_ were also accurate in most scenarios, including those where 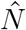 was biased. Note that confidence intervals of *s*_*l*_ were quite large, which is not surprising given that they are locus specific: they are thus based on a single trajectory (in contrast to *N*).

**Table 1.** Estimation of N and s for datasets simulated with a proportion *p* = 0.01 of loci under selection. We generated 100 replicates of datasets with 1000 loci and *n* = 30 at each of 10 sampling times *t* = 10, …, 100. Each row shows the estimations means and the median and coverage of 95% confidence intervals for: *N* (1^st^ row), *s*_*ℓ*_ of loci under selection (2^nd^ row) and 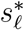 of neutral loci (3^rd^ row).

| | | $N = 100$ | | $N = 1000$ | |
| --- | --- | --- | --- | --- | --- |
| | | $s = 0.01$ | $s = 0.1$ | $s = 0.01$ | $s = 0.1$ |
| $\hat{N}$ | mean MLE | 98.35 | 98.06 | 1015.78 | 938.48 |
|  | median CI | 96 — 102 | 96 — 102 | 932 — 1106 | 866 — 1015 |
|  | CI coverage | 86% | 81% | 91% | 66% |
| $\hat{s}_\ell$ | mean MLE | 0.0231 | 0.2018 | 0.0106 | 0.1059 |
|  | median CI | -0.076 — 0.112 | -0.034 — 0.373 | -0.017 — 0.035 | 0.044 — 0.187 |
|  | CI coverage | 88.1% | 92.7% | 91.3% | 93.5% |
| $\hat{s}_\ell^*$ | mean MLE | -0.0003 | -0.0003 | 0 | 0 |
|  | median CI | -0.092 — 0.102 | -0.092 — 0.102 | -0.026 — 0.026 | -0.026 — 0.026 |
|  | CI coverage | 90.6% | 90.7% | 94.8% | 95.4% |

To investigate whether these good properties would still hold in more challenging scenarios, we next considered increasing proportions of loci under selection, up to the extreme case where all simulated loci are under selection. This generally led to a negative bias in *N* estimations, which increased with selection intensity and the proportion of the genome under selection (Figure 3). The effect was also more pronounced for larger populations, for instance in the worst case where all loci were selected with *s* = 0.1 the relative bias was about 25% for *N* = 100 and 75% for *N* = 1000. This last observation likely results from the fact that selection is more efficient in large populations; to phrase it in a different way, the effect of selection on genetic diversity is known to be better captured by the product *Ns* than by *s* alone (Paris et al., 2019a).

**Figure 3.**
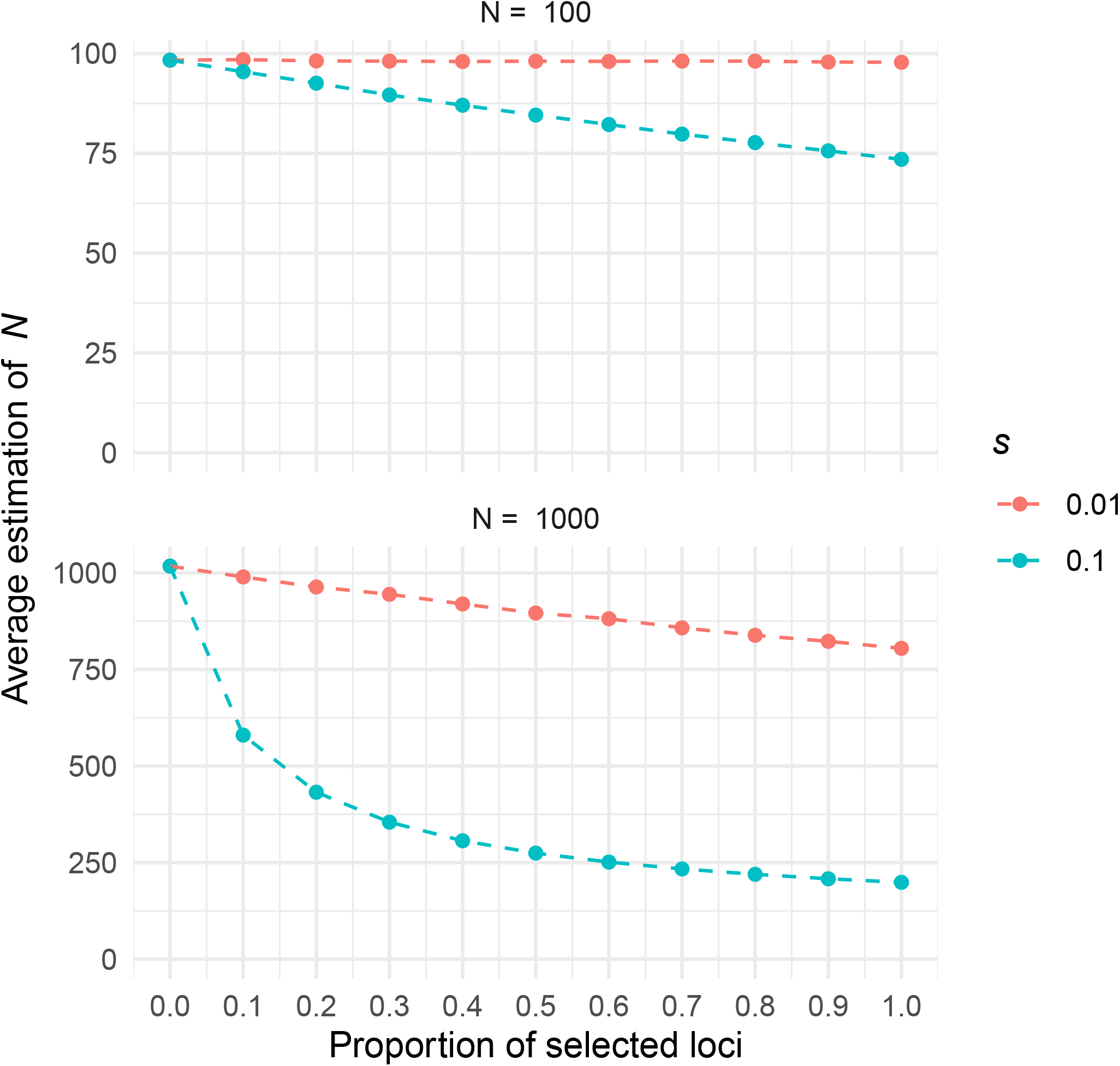
Estimation of the effective population size using SelNeTime, for simulated datasets with various proportions of loci actually under selection (x axis). For a given proportion *p*, 100 datasets of 1000 * *p* selected loci and (1 − *p*) * 1000 neutral loci were considered. Allele frequency trajectories at each locus were simulated independently under the Wright-Fisher model, with *n* = 30 at each of 10 sampling times *t* = 10, …, 100. Initial allele frequencies were set to *x*_0_ = 0.5 and the true population size was *N* = 100 (top) or *N* = 1000 (bottom). The average estimated value over the 100 datasets is shown on the y-axis, for different selection intensities (colors)

Interestingly, point estimations of *s*_*l*_ were not affected by this (potentially strong) bias in *N*, even in the most extreme case where all loci were under selection (Table S1, Figure S3). Only confidence intervals of *s* were affected and appeared to be larger when the negative bias of *N* was more pronounced. These properties directly result from the fact that *s*_*l*_ is related to the trend of the allele frequency trajectory, while *N* is related to its variance. An intuitive way to think about it is that providing to SelNeTime a too low value of *N* does not prevent it to capture the correct trend of the trajectory, but leads it to over-estimate its expected variance and to return a too large confidence interval for this trend.

Having established that the main issue for practical applications of SelNeTime concerns the estimation of *N* in the presence of selected sites, we finally studied whether this bias could potentially be avoided by accounting for the presence of selection. To do so, we estimated *N* on datasets where all loci were simulated under selection with the same intensity *s*_*T*_, but this time providing this true value to SelNeTime (i.e. computing the likelihood given *s* = *s*_*T*_ instead of *s* = 0). Results were much more accurate, as shown in Figure S4. Mean estimated values with and without the *s* = 0 assumption are summarized in Table 2.

**Table 2.** Mean estimated values of *N* over 100 replicates, for datasets simulated with different combinations of *N* and *s*_*T*_ and a sampling time interval Δ*t* = 10. Estimations assuming either *s* = 0 or *s* = *s*_*T*_ are compared.

| | $N_T = 100$<br>$s = 0$ | $N_T = 100$<br>$s = s_T$ | $N_T = 1000$<br>$s = 0$ | $N_T = 1000$<br>$s = s_T$ |
| --- | --- | --- | --- | --- |
| $s_T = 0$ | 98.4 | — | 1018 | — |
| $s_T = 0.01$ | 97.8 | 98.4 | 805 | 1014 |
| $s_T = 0.1$ | 73.5 | 98.5 | 199 | 1002 |

### Improved computation time

One important improvement of SelNeTime compared to the inital implementation of the BwS approach in compareHMM (Paris et al., 2019b) concerns the computation time, which was reduced using two main actions. First, the continuous density of the distribution is discretized when building the transition matrices (with a controlled bin width), allowing this continuous part of the distribution to be analyzed exactly in the same way as the discrete probability mass points in 0 and 1 (see section ‘The Beta with Spikes approximation‘). Second, matrix computations performed at each iteration of the HMM forward algorithm were vectorized, taking advantage of the Numpy library (Harris et al., 2020) to efficiently handle different loci or parameter values in a single operation.

As an illustration, Table 3 shows the computation time required to analyze a time series of 10 samples of 30 observations each, with a sampling time interval of Δ*t* = 10 generations and a number of loci varying from 100 to 10000.

**Table 3.** Computation times of CompareHMM (Paris et al., 2019b) and SelNeTime, run on a single CPU of an Intel Xeon 16 *\** 2.60 GHz processor with 32Gb of RAM.

| Nb. loci | compareHMM,<br>estimation of $s$ | SelNeTime, estimation<br>of $s$ | SelNeTime, estimation<br>of $N$ |
| --- | --- | --- | --- |
| 100 | 39.27s | 28.7s | 6.12s |
| 1000 | 360.01s | 36.21s | 14.23s |
| 10000 | 3530.47s | 96.95s | 87.83s |

For 100 loci, SelNeTime runs in approximately the same time as compareHMM, but the relative performance of SelNeTime improves quickly when the number of loci increases. Actually, the time to compute all transition matrices, which is the first step of both SelNeTime and compareHMM analyses, is independent of the number of loci (but it depends on the sampling design). This part of the analysis took approximately 28s for SelNeTime in this example, which was of the same order as compareHMM, but after this the time spent per locus was far lower with SelNeTime (approx. 0.007s) than with compareHMM (approx. 0.35s).

### Comparison with the Wright-Fisher model

While SelNeTime computes by default HMM transitions using the BwS distribution, it might occasionally be useful to compute them based on the WF model, in particular to evaluate the impact of the BwS approximation on inference quality. Although this is numerically possible only for small population sizes (not more than a few hundreds), we included one option to do this in the program. For a true population size of *N* = 100, we could for instance check that using the BwS or the WF model had very limited impact on the inference of either effective population size or selection (Figure S5). A more detailed comparison between the estimations obtained by the BwS and WF distributions can be found in Paris et al. (2019a), but only for selection inference.

### Simulation of datasets

As already observed in Paris et al. (2019a) and several other studies on the subject, the estimation accuracy of effective population size or selection depends on both the true evolutionary parameters of the population under study (*N* and *s*) and the sampling design of observed data (number and size of observed samples, time intervals between samples). In order to help researchers plan future experiments, or interpret the results obtained for a specific dataset, SelNeTime includes a simulator of observed allele frequency trajectories. Simulations can easily be tunned using either a yaml file or command line options and output data at a format that can directly be analyzed with other SelNeTime commands. Notably, this simulator was used to generate the data underlying the figures and tables of this study.

## Discussion

The SelNeTime python package provides a re-implementation of the BwS approach proposed by Paris et al. (2019a) to analyse genomic time series data. Compared to the initial implementation of these authors (Paris et al., 2019b), SelNeTime (i) estimates the effective population size during the temporal period covered by the samples, (ii) runs much faster, epecially for large SNP datasets, and (iii) provides a user-friendly interface with simple commands, including a simulator.

In a first set of simulation scenarios, we validated the fact that SelNeTime accurately infers effective population size in the absence of selection, or selection intensity at each locus assuming the true value of *N* is provided. Obviously, these two hypothetical conditions are almost never met in practice: populations are subject to selection constraints and their effective population size is generally unknown. However, what is much less clear is the proportion of the genome that actually evolves as under selection. On one hand, because a large proportion of the genome is non functional and even functional sites do not necessarily impact fitness related traits, one might expect this proportion to be small. On the other hand, accounting for negative (rather than only adaptive/positive) selection and for indirect selection effects due to genetic linkage might substantially increase this proportion and recent empirical studies suggested that linked selection would actually be pervasive in the genome of species like humans (Pouyet et al. (2018)) or drosophila (Elyashiv et al. (2016)). In a second set of simulation scenarios, we thus explored the performance of SelNeTime when both *N* and the *s*_*l*_ must be estimated, considering various proportions of the genome under selection.

If selection is relatively rare in the genome, SelNeTime was found to accurately infer both *N* and the *s*_*l*_ (Table 1). For higher proportions of selected sites, *N* was always under-estimated, but the extent of this bias depended on the true values of both *N* and *s* and was for instance limited for small populations or weak selection (Figure 3). Importantly, selection intensity was always accurately inferred, whatever the bias observed for 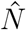 (Table S1). This results is consistent with several recent studies using genomic time series, which suggested that accurate estimations of selection could be achieved despite demographic misspecification (Mathieson & Terhorst, 2022; Whitehouse & Schrider, 2023). Overall, these results suggest that SelNeTime will provide a good estimation of the parameters of interest in a large set of practical situations. Indeed, the original compareHMM approach (Paris et al., 2019b), which is very close to SelNeTime for the estimation of selection parameters, has already been applied to several empirical datasets and provided relevant biological insights (Paris et al., 2019a; Boitard et al., 2021, 2023).

Because the estimation of *N* might still be an issue when a large part of the genome is affected by selection, a main objective of future research will be to extend the current approach in order to really co-estimate demography and selection. While this is clearly a challenging task, which very few population genetics studies tried to tackle so far, we point out that SelNeTime estimation of *N* is also accurate in the presence of selection, provided the true value of selection intensity is given as input (Figure S4). Encouragingly, this result suggests that optimization approaches accounting for the effects of both *N* and the *s*_*l*_’s on the likelihood, could perform well.

Another assumption made by SelNeTime is to consider that allele frequency trajectories observed at distinct loci are independent, an assumption that does not hold in practice for at least two reasons: (i) linkage disequilibrium (LD) and (ii) the so-called Bulmer effect (Bulmer, 1971), i.e. the negative correlation of allele frequencies resulting from the polygenic selection of quantitative traits. While the LD effect can easily be mitigated by sub-sampling SNPs, the Bulmer effect would be more tricky to model with our approach. One first step to explore this question could be to apply SelNeTime to genomic time series simulated under polygenic selection scenarios. In the longer term, we could also try to estimate temporal variations of *s*_*l*_ at each locus (see Mathieson & Terhorst (2022)), which would be more consistent with a model of selection on quantitative traits. Importantly, based on theoretical properties of composite likelihoods, we do not expect that SelNeTime estimates of *N* will be biased in the presence of correlations between loci; only the confidence intervals, such as those shown in Figure S5 and returned in the output of the software, may be affected.

Finally, admixture might be another important counfouding effect for our approach, as recently outlined for another method analyzing genomic time series under the assumption of panmixia (Simon & Coop, 2024). The impact of admixture on SelNeTime performance will likely depend on the exact admixture scenario (admixture proportions, single versus recurrent events, overlap of these events with the sampling period, divergence with the source population, local versus global adaptation …) so a first step could again be to evaluate it using simulations. It should also be feasible to modify SelNeTime HMM transitions in order to account for specific well documented admixture scenarios, in a similar spirit of what was done in Simon & Coop (2024).

## Data, script and code availability

SelNeTime is available on the python package index https://pypi.org/project/selnetime and on the project git repository https://forge.inrae.fr/genetic-time-series/selnetime. All figures of this study can be reproduced using a snakemake pipeline available at https://forge.inrae.fr/genetic-time-series/snt-simulation-pipeline.

## Acknowledgments

A preprint version of this article has been peer-reviewed and recommended by PCI Math Comp Biol (https://doi.org/10.24072/pci.mcb.100406).

We would like to thank Alan Rogers (recommender), Rui Borges (reviewer) and one anonymous reviewer for their constructive comments on the manuscript.

## Supplementary material

**Figure S1.**
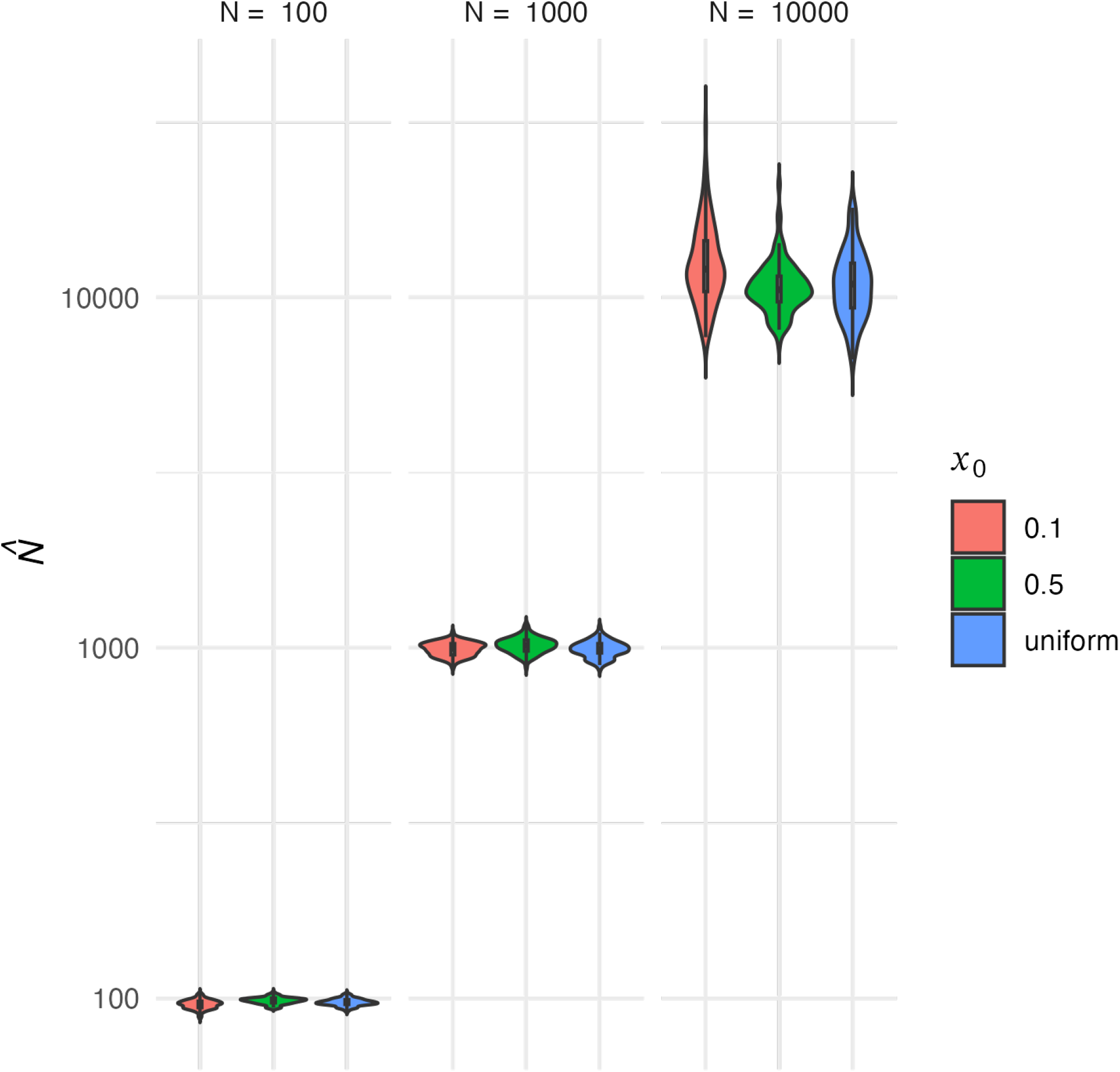
Estimation of the effective population size using SelNeTime, for different values of *N* (panels) and different values / distributions of the initial allele frequency *x*_0_ (colors). For each combination of these two parameters, 100 datasets of 1000 loci were considered. Allele frequency trajectories at each locus were simulated independently under the Wright-Fisher model, with *n* = 30 at each of 10 sampling times *t* = 10, …, 100.

**Figure S2.**
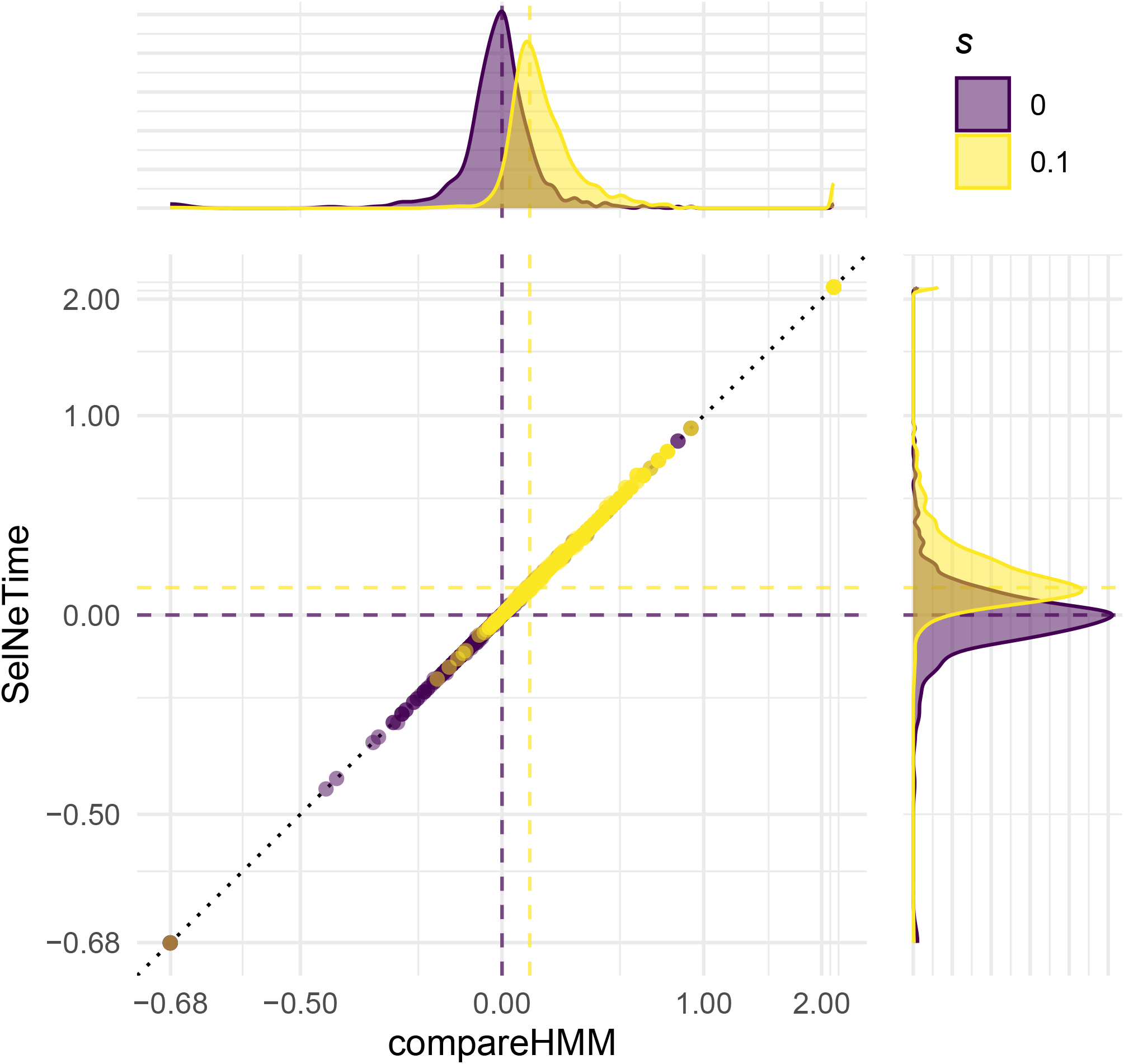
Estimation of the selection intensity for 1000 loci using SelNeTime and compareHMM. Allele frequency trajectories at each locus were simulated independently under the Wright-Fisher model with *N* = 100, *s* = 0 (purple) or *s* = 0.1 (yellow) and an initial allele frequency *x*_0_ = 0.5. Allele frequencies were observed for *n* = 30 at 10 sampling times *t* = 10 … 100. A log_2_(1 + *x*) transformation was applied to both axes, in order to have a qualitatively symmetrical scale around 0 (a selection of *−*0.5 is the opposite of *s* = 1, etc.)

**Figure S3.**
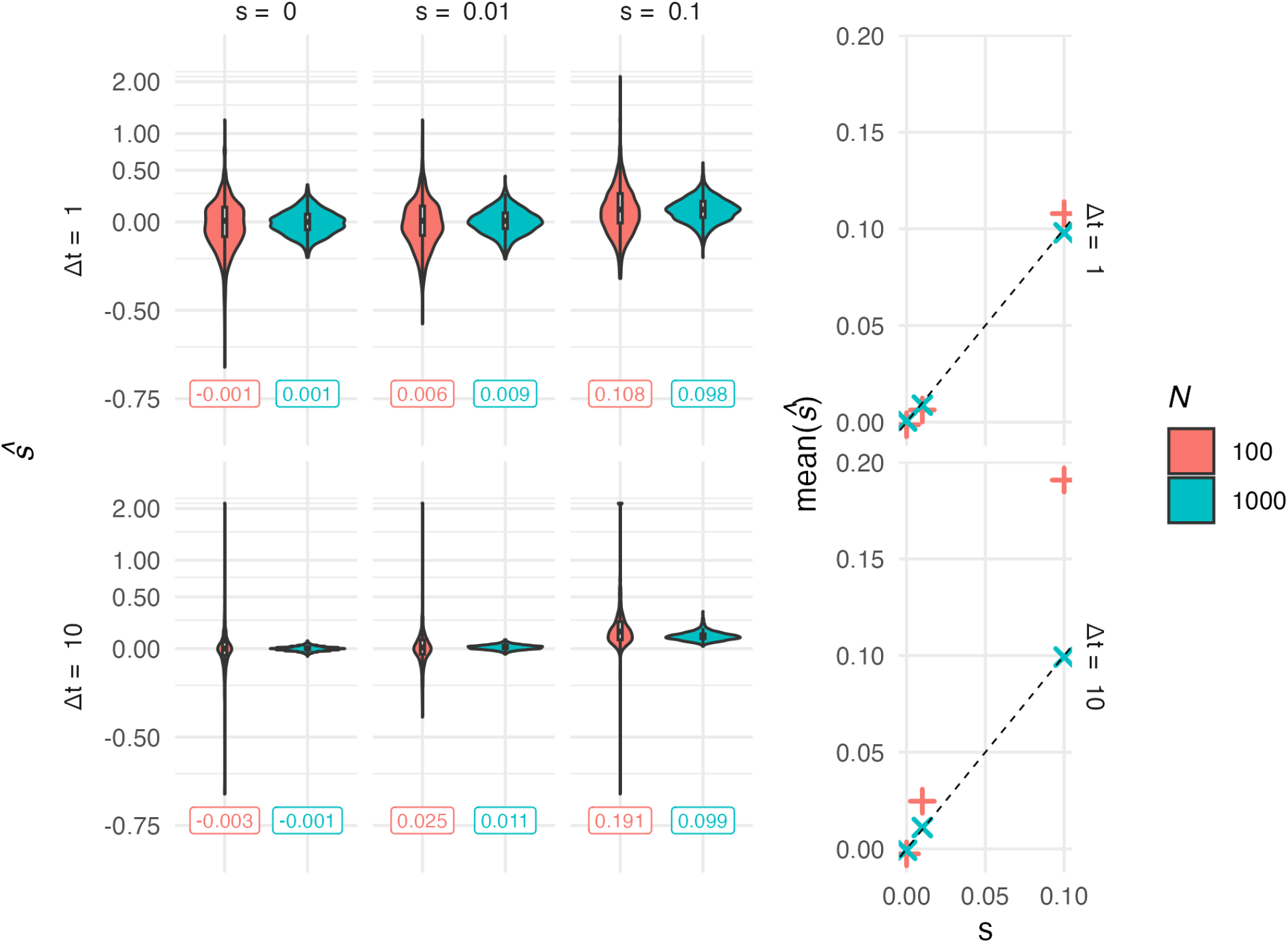
Estimation of the selection intensity using SelNeTime without knowing *N*. For given values of *s* (columns) and *N* (colors), 1000 loci were considered. Allele frequency trajectories at each locus were simulated independently under the Wright-Fisher model, with *n* = 30 at each of 10 sampling times *t* = 1, …, 10 or *t* = 10, …, 100 (lines), and an initial allele frequency *x*_0_ = 0.5. Distribution of estimations for the 100 loci. The mean value for each simulation scenario is shown below the associated violin plot, and on the right part of the plot. A log_2_(1 + *x*) transformation was applied to the y axis of the left panel.

**Figure S4.**
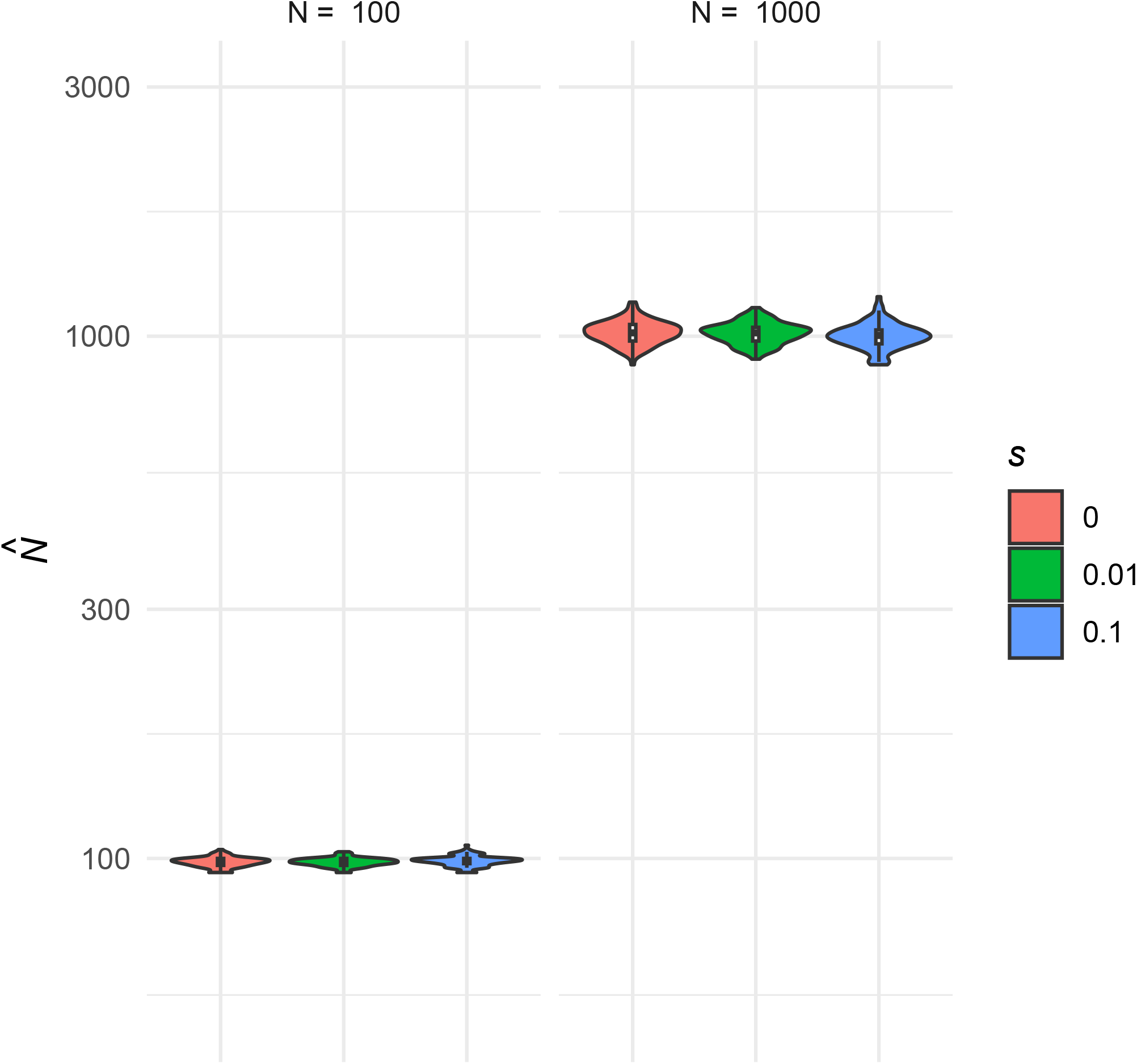
Estimation of the effective population size using SelNeTime with the true value of *s* provided, for simulated datasets where all loci are under selection with the same intensity. For given values of *s* (colors) and *N* (panels) 100 datasets of 1000 loci were considered. Allele frequency trajectories at each locus were simulated independently under the Wright-Fisher model, with *n* = 30 at each of 10 sampling times *t* = 10, …, 100. Initial allele frequencies were set to *x*_0_ = 0.5.

**Figure S5.**
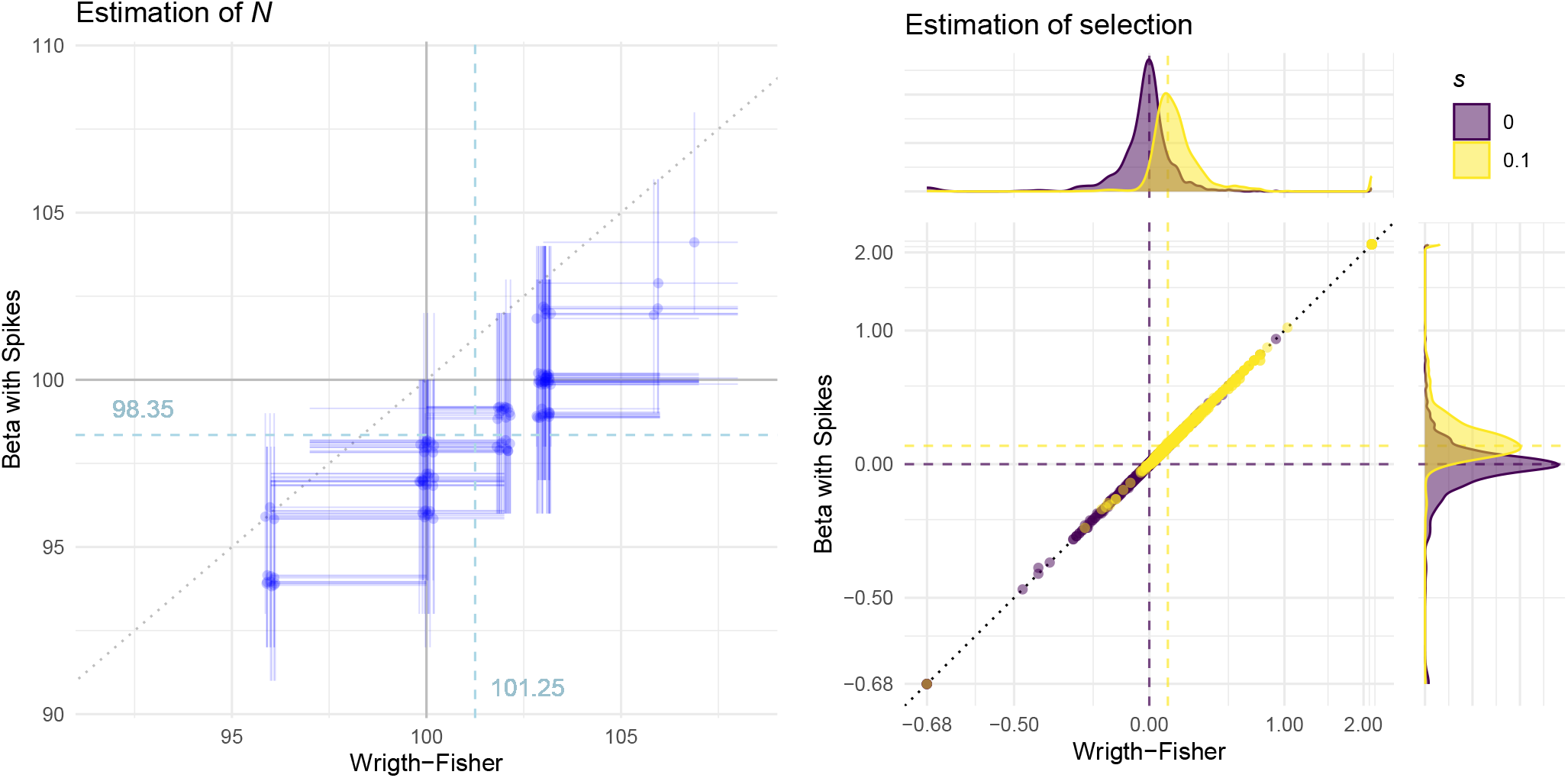
Comparison of SelNeTime estimations based on the Beta with Spikes and the Wright-Fisher model. **(a)**: MLE and 95% confidence interval of effective population size for 100 datasets of 1000 loci each. **(b)** MLE of *s* for 1000 loci. The two panels show the estimations obtained using the WF (x axis) or the BwS (y axis) model on the same data, which were simulated independently under the Wright-Fisher model with *N* = 100 *s* = 0 (Left and Right, purple) or *s* = 0.1 (Right, yellow) and initial allele frequencies *x*_0_ = 0.5. Allele frequencies were observed for *n* = 30 at 10 sampling times *t* = 10…100. A log_2_(1 + *x*) transformation was applied to both axes of panel b in order to restore the symmetry between positive and negative values of *s* (for instance, *s* = −0.5 implies that the fitness of reference homozygote genotypes is twice that of alternative homozygote genotypes, the symmetric case of *s* = 1).

**Table S1.** Estimation of N and s for datasets simulated with all loci under selection. We generated 100 replicates of datasets with 1000 loci and *n* = 30 at each of 10 sampling times *t* = 10, …, 100. Each row shows the estimations mean and the median and coverage of 95% confidence intervals for: *N* (1^st^ row), and *s*_*ℓ*_ of loci under selection (2^nd^ row).

| | | $N = 100$ | | $N = 1000$ | |
| --- | --- | --- | --- | --- | --- |
| | | $s = 0.01$ | $s = 0.1$ | $s = 0.01$ | $s = 0.1$ |
| $\hat{N}$ | mean MLE | 97.81 | 73.5 | 804.77 | 198.96 |
|  | median CI | 96 — 100 | 71 — 76 | 743 — 860 | 192 — 206 |
|  | CI coverage | 82% | 0% | 0% | 0% |
| $\hat{s}_\ell$ | mean MLE | 0.0220 | 0.1978 | 0.0109 | 0.0995 |
|  | median CI | -0.076 — 0.122 | -0.059 — 0.411 | -0.017 — 0.044 | -0.009 — 0.236 |
|  | CI coverage | 89.5% | 95.9% | 93.5% | 99.8% |

